# Integration of gut microbiome, host biomarkers, and urine metabolome data reveals networks of interactions associated with distinct clinical phenotypes

**DOI:** 10.1101/509703

**Authors:** Rui-Jun Li, Zhu-Ye Jie, Qiang Feng, Rui-Ling Fang, Fei Li, Yuan Gao, Hui-Hua Xia, Huan-Zi Zhong, Bin Tong, Jia-Yu Xu, Lise Madsen, Chun-Lei Liu, Zhen-Guo Xu, Jian Wang, Huan-Ming Yang, Xun Xu, Yong Hou, Susanne Brix, Karsten Kristiansen, RXin-Lei Yu, Hui-Jue Jia, Kun-Lun He

## Abstract

**Background:** Comprehensive analyses of multi-omics data may provide insights into interactions between different biological layers in relation to distinct clinical features. Here we integrated data on the gut microbiota, blood parameters and urine metabolites of treatment-naive individuals presenting a wide range of metabolic disease phenotypes to delineate clinically meaningful associations.

**Results:** Trans-omics correlation networks revealed that candidate gut microbial biomarkers and urine metabolite features covaried with distinct clinical phenotypes. Integration of the gut microbiome, the urine metabolome and the phenome revealed that variations in one of these three systems correlated with changes in the other two. Of specific note in relation to clinical parameters of liver function, we identified *Eubacterium eligens, Faecalibacterium prausnitzii* and *Ruminococcus lactaris* to be associated with a healthy liver function, whereas *Clostridium bolteae*, *Tyzzerella nexills, Ruminococcus gnavus, Blautia hansenii*, and *Atopobium parvulum* were associated with blood biomarkers for liver diseases. Variations in these microbiota features paralleled changes in specific urine metabolites. Network modeling yielded two core clusters including one large gut microbe-urine metabolite close-knit cluster and one triangular cluster composed of a gut microbe-blood-urine network, demonstrating close inter-system crosstalk especially between the gut microbiome and the urine metabolome.

**Conclusions:** Distinct clinical phenotypes are manifested in both the gut microbiome and the urine metabolome, and inter-domain connectivity takes the form of high-dimensional networks. Such networks may further our understanding of complex biological systems, and may provide a basis for identifying biomarkers for diseases.

## Background

Systemic metabolism is regulated at numerous levels. Apart from direct interactions, the gut microbiota may indirectly interact with the host via metabolites, which may appear in circulation and the urine. Considering the complexity and the inter-dependency of biological systems, a holistic approach integrating multiple omics data may further the understanding of metabolic disease development.

The gut microbiota is a metabolic “organ” co-evolving with the host. Composed of hundreds of trillions of microbes [1], it is an ecological community where members compete, cooperate, and synergize with each other. This community not only passively adapts to the local biotic microenvironment, but also actively interacts with the host. The gut microbiota affects the integrity of the gut barrier, extracts energy from food, provides bioactive compounds, regulates metabolic homeostasis and acts as a pro-inflammatory-anti-inflammatory rheostat [2]. The composition and functional capacity of the gut microbiota may hence reflect the physiological state of the host, providing information of possible pathological conditions. This notion is supported by results from several case-control studies reporting on gut microbiota signatures that are linked to metabolic diseases, such as obesity and diabetes [3-5]. Perturbations of the microbiota may cause alterations in the metabolite profile in circulation [4, 6, 7], and in addition, the compositional and functional features of the gut microbiota may also be detectable in the urine [8]. Combining data on the composition and functional potential of the gut microbiome with blood and urine metabolite profiles may reveal novel associations with clinical phenotypes and thereby provide new knowledge of host-microbiome interactions in health and disease.

Taking advantage of a cohort of treatment-naïve subjects representing various clinical phenotypes, we performed an integrative trans-omics study combining blood biomarkers, fecal metagenomics and urine metabolomics. Correlation analysis between the blood profile and one of the other two omics datasets identified potential biomarkers characterizing different phenotypes, while integration of the three omics datasets revealed novel covariations and links. Community modeling further revealed the intimate interaction among the three systems, especially between the gut microbiota and the urine metabolome.

## Results

### Characteristics of the study cohort and clinical parameters

We included 138 treatment-naïve subjects who presented wide variations in clinical blood parameters (based on 130 blood biomarkers, including biomarkers associated with metabolic abnormalities such as hyperglycemia, hyperinsulinemia, hyperlipidemia, and kidney dysfunction) (Additional file 1: Table S1; Additional file 2: Table S2; Additional file 3: Table S3). Amongst these treatment-naïve subjects, we have an overrepresentation of individuals diagnosed with carotid atherosclerosis at the time of sampling (n=102), as the study subjects took part in a specifically designed screening program for carotid atherosclerosis. Due to their display of a diverse range of metabolic abnormalities and a large variation in blood biomarkers, the collected data allowed us to evaluate possible associations between gut microbiota composition, clinical parameters and urine metabolites across different clinical states.

### Selective clinical parameters associate with the gut microbiota profile

Shotgun-based metagenomic sequencing of fecal microbial DNA generated a total of 1110.22 Gb of data (on average 7.93 Gb data per sample), which we clustered into 645 Metagenomic species (MGS) [9] present in more than 10% of samples (Additional file 4: Table S4).

To examine associations between phenotypes and the overall microbiota profile, we performed Permutational multivariate analysis of variance (PERMANOVA) of each variable vs MGS compositions. Thirty two variables were identified as covariates, of which gamma-glutamyl transferase (GGT, r^2^=0.022), ferritin (r^2^=0.020), alanine aminotransferase (ALT, r^2^=0.020), monocytes (r^2^=0.020), and triglyceride (TRIG, r^2^=0.018) had the largest effect sizes (p<0.05, q<0.1, Fig. 1A). Five clinical covariates correlated significantly with α-diversity, and 6 clinical covariates correlated significantly with total gene counts in the gut microbiota (Additional file 5: Fig S1). Conversely, we also tested whether the microbiome data could be used to predict phenotypic variations. We built prediction models using Random forest based on the relative abundance of MGSs. The gut microbiota-based predictor performed best in the prediction of TRIG (r=0.35), followed by low fluorescence reticulocytes (LFR, r=0.28), middle fluorescent reticulocytes (MFR, r=0.25), hematocrit (r=0.21), and uric acid (URIC, r=0.21) (Fig. 1B, and Additional file 6: Fig S2), indicating generally weak associations between certain blood biomarkers and the overall composition of the gut microbiota.

**Fig. 1.**
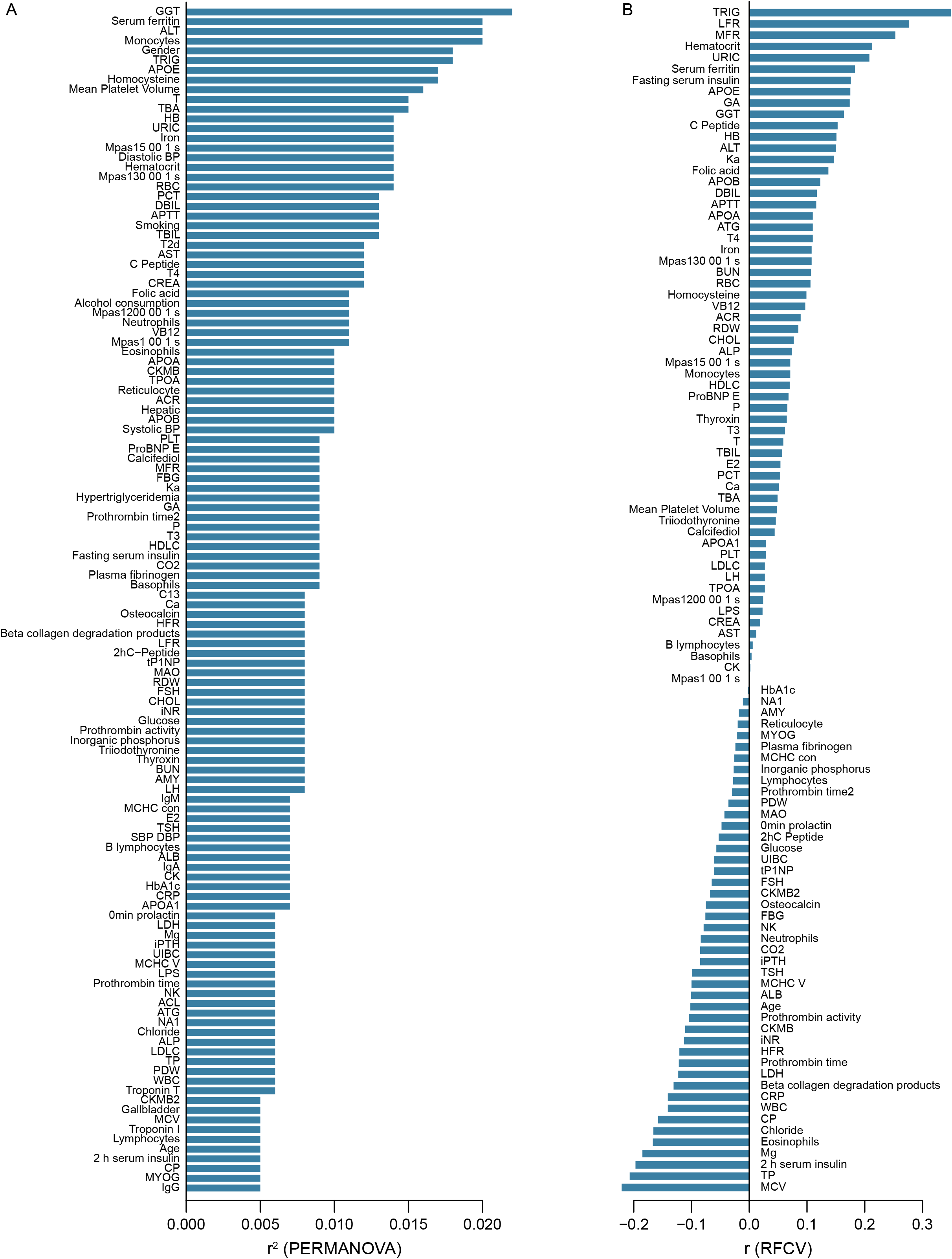
Alterations of the gut microbiome community associated with clinical parameters. **(a)** Clinical parameters that showed associations with MGS composition in PERMANOVA, ranked by the effect size (r^2^). p<0.05, q<0.1 **(b)** Prediction of clinical parameters by MGS relative abundances with random forest models, ranked by the performance (the correlation coefficient between predicted value and the measured value). p<0.05.

### Microbial taxa and functions correlated with clinical phenotypes

We next aimed to identify potential biomarkers in the gut microbiome that could be linked to clinical parameters. Spearman correlation analysis was conducted between individual MGSs and clinical parameters. Among the FDR-adjusted significant correlations, the abundance of *Eubacterium eligens* correlated negatively with GGT (MGS 0507, r=-0.30; MGS1459, r=-0.31; MGS1432, r=-0.33) and total blood levels bilirubin (TBIL) (MGS1459, r=-0.26; MGS1432, r=-0.28; MGS 0507, r=-0.33;), whereas the abundances of *Tyzzerella nexilis* (MGS0415, r=0.30) and *Blautia hansenii* (MGS0787, r=0.30) correlated positively with GGT. *Atopobium parvulum* (MGS1482, r=0.39) and *Solobacterium moorei* (MGS0945, r=0.35) correlated positively with blood levels of thyroxine (T4) (p<0.05, q<0.1, Fig. 2, and Additional file 7: Table S5). In line with this, *Eubacterium eligens* and *Atopobium parvulum* have been reported to be depleted and enriched, respectively, in patients with atherosclerotic cardiovascular disease [10].

**Fig. 2.**
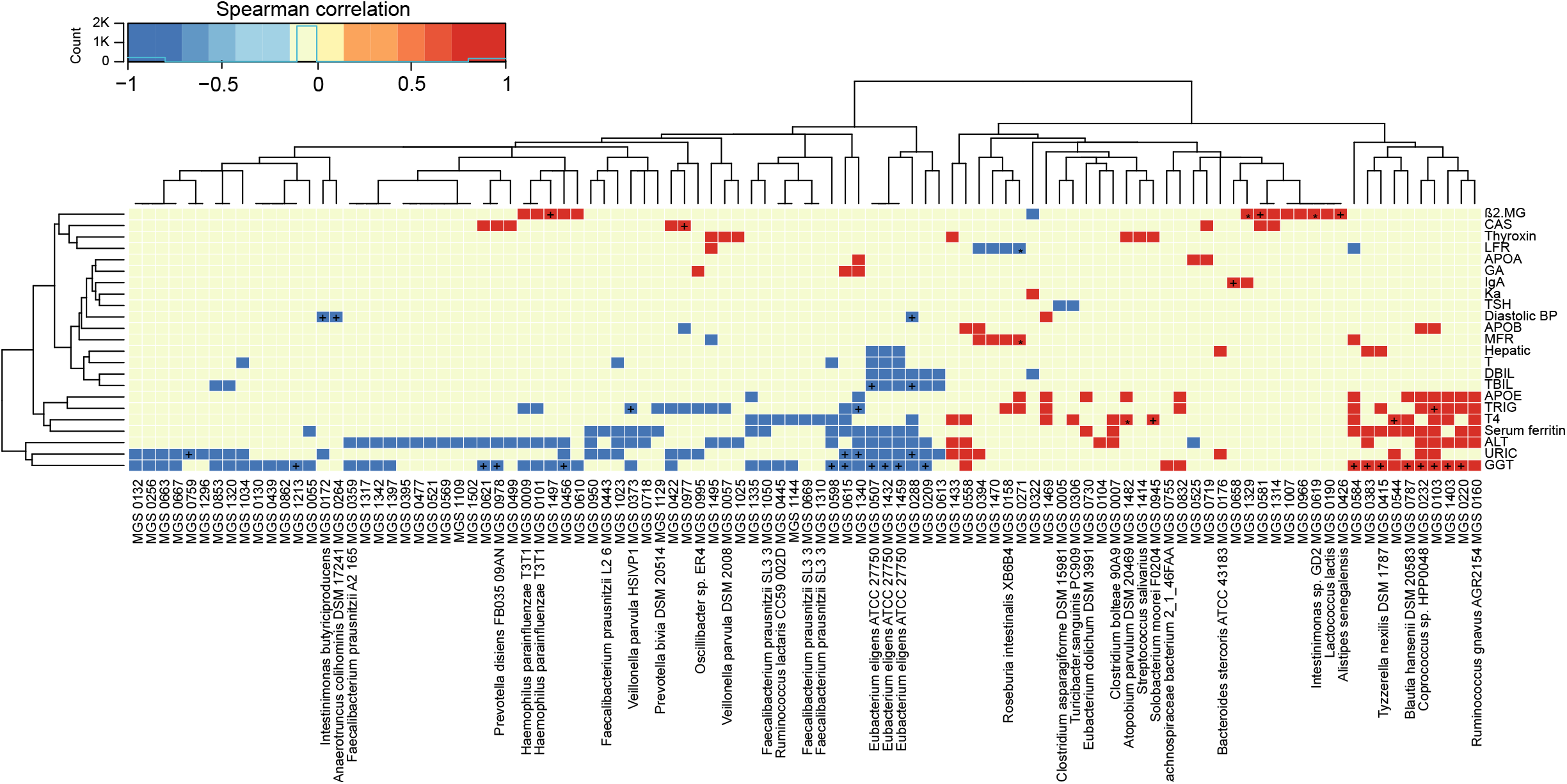
Correlations between the gut microbiome and clinical parameters. Heatmap showing correlations between MGS and clinical parameters (Spearman correlation, p<0.05). +, q<0.1; *, q<0.01. Color scale indicates the value of correlation coefficient.

Next, we analyzed the correlation between microbial functions and clinical parameters. The abundances of genes encoding proteins involved in amino acid metabolism (Proline biosynthesis, M00015, r=-0.27; Cysteine biosynthesis, M00021, r=-0.32; Tryptophan biosynthesis, M00023, r=-0.26; Histidine biosynthesis, M00026, r=-0.30; Ornithine biosynthesis, M00028, r=-0.31; Methionine biosynthesis, M00017, r=-0.28), gene expression (Aminoacyl–tRNA biosynthesis, M00359, r=-0.29; Aminoacyl-tRNA biosynthesis, M00360, r=-0.27; RNA polymerase, M00183, r=-0.30; Ribosome, M00178, r=-0.29; Ribosome, M00179, r=-0.30), and energy production (Coenzyme A biosynthesis, M00120, r=-0.27; NAD biosynthesis, M00115, r=-0.32; Pantothenate biosynthesis, M00119, r=-0.27) correlated negatively with monocyte counts, and the abundances of genes encoding proteins involved in tyrosine degradation (M00044, r=-0.31), and teichoic acid transport system (M00251,r=-0.30) correlated negatively with basophils. The abundances of genes involved in the transport of nickel and/or cobalt (M00246, r=0.37; M00245, r=0.36) positively correlated with iron (Additional file 8: Fig S3; and Additional file 9: Table S6).

These correlations indicated that alterations in the microbiota taxa and function may be linked to specific physiological perturbations.

### Urine metabolomic variations associated with clinical phenotypes

The metabolites in the urine may be generated by the host and /or the gut microbiota. By metabolome profiling of the urine, we detected 10890 positively charged and 9416 negatively charged metabolites, which were clustered into 1042 modules (more than 3 metabolites in each module) based on their co-abundance using the R package WGCNA [6]. To assess their clinical relevance, we performed correlation analysis between urine metabolites and clinical parameters. A large number of urine metabolites correlated with blood biomarkers, compartmentalized into distinct clusters exhibiting positive or negative correlations (Fig. 3; Additional file 10: Fig S4; Additional file 11: Table S7). These urine metabolites demonstrated opposing correlation relationships with two sets of blood biomarkers. The metabolites that correlated positively with one set of blood biomarkers including follicle-stimulating hormone (FSH), luteotropic hormone (LH), glycated albumin (GA), brain natriuretic peptide brain natriuretic precursur (proBNP), high-density lipoprotein cholesterol (HDLC), vitamin B12 (VB12), calcifediol, beta-collagen degradation products, osteocalcin, total propeptide of type I procollagen (tP1NP), fibrinogen, and alkaline phosphatase (ALP), tended to correlate negatively with blood parameters related to liver function (URIC, creatinine [CREA], ALT, GGT, AST, TBIL, and direct bilirubin [DBIL]), blood pressure, lipid metabolism (apolipoprotein B [APOB], apolipoprotein E [APOE], TRIG), and glucose metabolism (C peptide, insulin, and glucose), and vice versa (Fig. 3, and Additional file 10: Fig S4). Among the phenotype-associated metabolites, those that correlated negatively with markers of liver dysfunction (e.g. URIC, CREA, ALT, GGT) included phenols, steroids, and possible food-derived compounds and metabolic intermediates (e.g. compounds structurally similar to fragomine, L–aspartyl–L–phenylalanine cycloheptanecarboxylic acid and, (R)–mevalonate, L–trans–5–hydroxy–2–piperidinecarboxylic acid, 20,26–dihydroxyecdysone), whereas metabolites that correlated positively included molecules that were possible metabolic intermediates or bioactive compounds (e.g. compounds structurally similar to ethyl cellulose, epomediol, 7–aminomethyl–7–carbaguanine, imipenem, S-adenosyl-L-homocysteine), suggesting the physiological involvement of those metabolites and related metabolic pathways. Thus, the urine metabolomic patterns reflected selective physiological features.

**Fig. 3.**
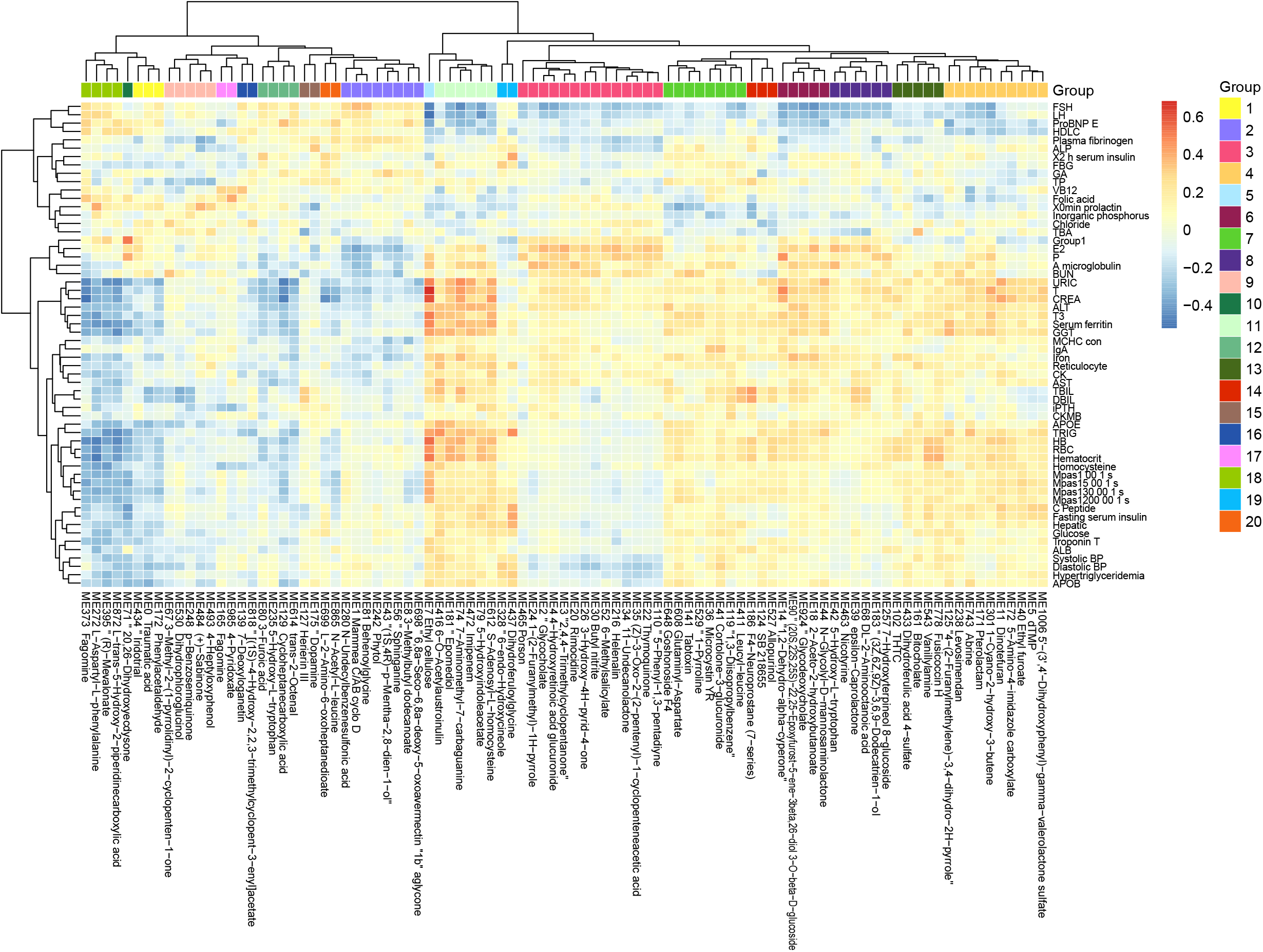
Correlations between the urine metabolome and clinical parameters. Heatmap showing correlations between urine metabolomic groups and clinical parameters (Spearman correlation, p<0.05, q<0.05). Only metabolites correlated significantly with more than one clinical parameter are included in the heatmap (the complete heatmap showing all correlations is illustrated in Additional file 10: Fig S4). Color scale indicates the value of correlation coefficient.

### Integration of the gut microbiome, the urine metabolome and the phenome

To integrate data on the gut microbiome, the urine metabolome and the phenome, we performed three rounds of pairwise Spearman correlations, retaining triangular correlations for p<0.05 (Fig. 4, and Additional file 12: Table S8). Compartmentalized by their direction of correlation with clinical parameters, we found microbe-metabolite pairs showing positive correlations (which we termed disease-related microbes/metabolites) and negative correlations (which we termed health-related microbes/metabolites) with blood biomarkers where elevated levels have been associated with diseases (Fig. 4). The correlation analyses indicated that for liver dysfunction-associated parameters (e.g. GGT and ALT), the disease-related bacteria included *Clostridium bolteae* (MGS0007), *Tyzzerella nexills* (MGS0415), *Ruminococcus gnavus* (MGS0160), *Blautia hansenii* (MGS0787), and *Atopobium parvulum* (MGS1482), whereas the health-related bacteria included *Eubacterium eligens* (MGS1432, MGS0507, and MGS1459), *Faecalibacterium prausnitzii* (MGS1310), and *Ruminococcus lactaris* (MGS0445). These bacteria shared a collection of correlated health-related or disease-related metabolites; the disease-related metabolites included molecules structurally similar to cinncassiol D2 glucoside, ethyl tiglate, and dihydroferulic acid 4-sulfate, and the health-related ones included those similar to phenylacetaldehyde, 3-methyl-quinolin-2-ol, 6-methylsalicylate, p-methoxycinnarnaldehyde, cassaidine, lithocholate 3-o-gluronide, and gentisate aldehyde (Fig. 4). Blood levels of APOE and/or TRIG correlated positively with the abundance of the disease-related bacteria *Ruminococcus gnavus, Blautia hansenii*, and *Atopobium parvulum*, which in turn correlated with a series of urine metabolites. These included the health-related metabolites structurally similar to phenylacetaldehyde, 3-methyl-quinolin-2-ol and lithocholate, and the disease-related metabolites similar to cinncassiol D2 glucoside and ethyl tiglate.

**Fig. 4.**
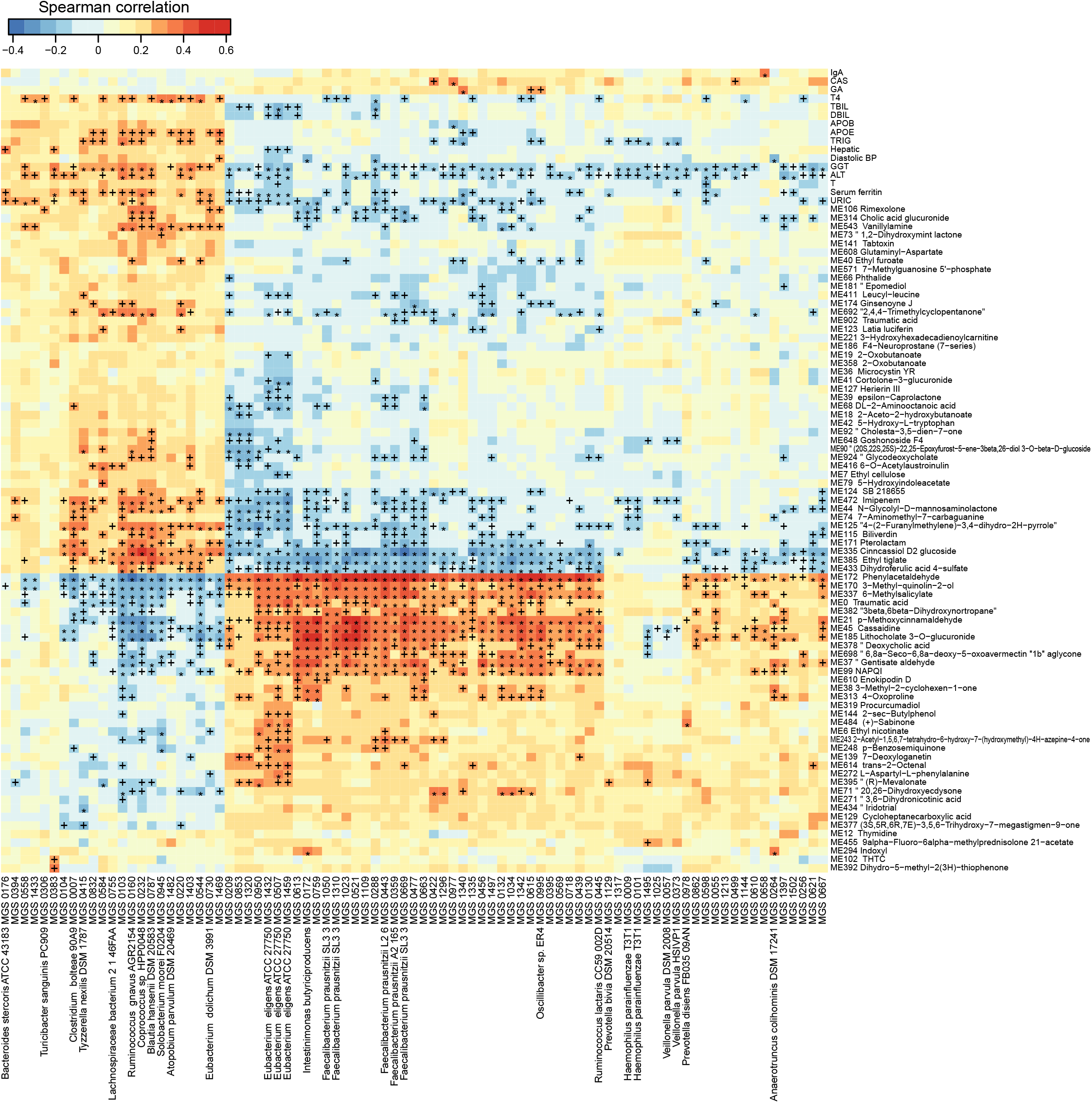
Trans-omics correlations between the gut microbiome, the urine metabolome and clinical parameters. Correlations that reached statistical significance (Spearman correlation, p<0.05) between MGS and clinical parameters, between metabolites and clinical parameters, and between MGS and metabolites, are shown in the heatmap. +, q<0.1; *, q<0.01. Color scale indicates the value of correlation coefficient.

Next, we applied the same inter-correlation approach to microbial modules, urine metabolites and blood biomarkers (p<0.05, q<0. 1; Additional file 13: Fig S5; Additional file 14: Table S9). A few modules significantly correlated with URIC (e.g. modules in carbohydrate transport and metabolism, M00171, M00169, M00172, M00491, M00194, M00200, M00206, and M00201), and these modules correlated with the health-related metabolites (e.g. metabolites structurally similar to lithocholate 3-o-gluronide, methoxycinnarnaldehyde, or phenylacetaldehyde) and the disease-related metabolites (e.g. metabolites structurally similar to ethyl tiglate, dihydroferulic acid or imipenem).

Taken together, results suggested that certain microbiome features, urine metabolites, and clinical parameters co-vary across treatment-naïve subjects.

### Organization of trans-domain correlations into one two-domain tight-knit and one three-domain inter-connected communities

Biological entities, such as microorganisms, exist in communities and interact with each other [11]. The interaction may be directly or indirectly via biomolecules. To unravel the structure of interactive relationship between the gut microbiota, urine metabolites and clinical parameters, we modeled correlations with a previously reported algorithm based on edge betweenness [11]. After pruning the loose connections, two large communities emerged (Fig. 5A). The largest community was densely weaved between 200 MGS and 33 metabolite groups (with 99.1%/1040 edges), partially linked to five blood biomarkers (with 0.2%/3 edges between MGS and phenotypes, and 0.7%/6 edges between metabolites and phenotypes, Fig. 5B, and Additional file 15: Table S10). This community represented a huge cluster of microbe-metabolite interactions, demonstrating extensive linkage between the gut microbiota and the urine metabolome. The second largest microbial community was formed by triangular correlations between 41 MGS, 53 metabolite groups and 23 blood biomarkers (with 5%/14 edges between MGS and phenotypes, 67%/181 edges between metabolites and phenotypes, and 28%/75 edges between MGS and metabolites), which was less dense than the first community (Fig. 5C). In this community, entities from the three systems were all connected, where changes in one data set resulted in variations in the other two. The majority of inter-system correlations were positive, except that a metabolite structurally similar to diethyl L–malate correlated negatively with blood variables including GGT, URIC, TRIG, blood viscosity, testosterone (T), triiodothyronine (T3), ferritin, hematocrit, monocytes, red blood cell count (RBC), hemoglobin (HB), gender, alcohol consumption and smoking, and with one MGS (MGS0544), and that a metabolite structurally similar to 5–amino–4–imidazole carboxylate correlated negatively with CREA, URIC and T, indicating antagonistic or inhibitive relationships. These modeled communities demonstrated close host-microbiome interactions and trans-omics connections, especially between the gut microbiota and the urine metabolome.

**Fig. 5.**
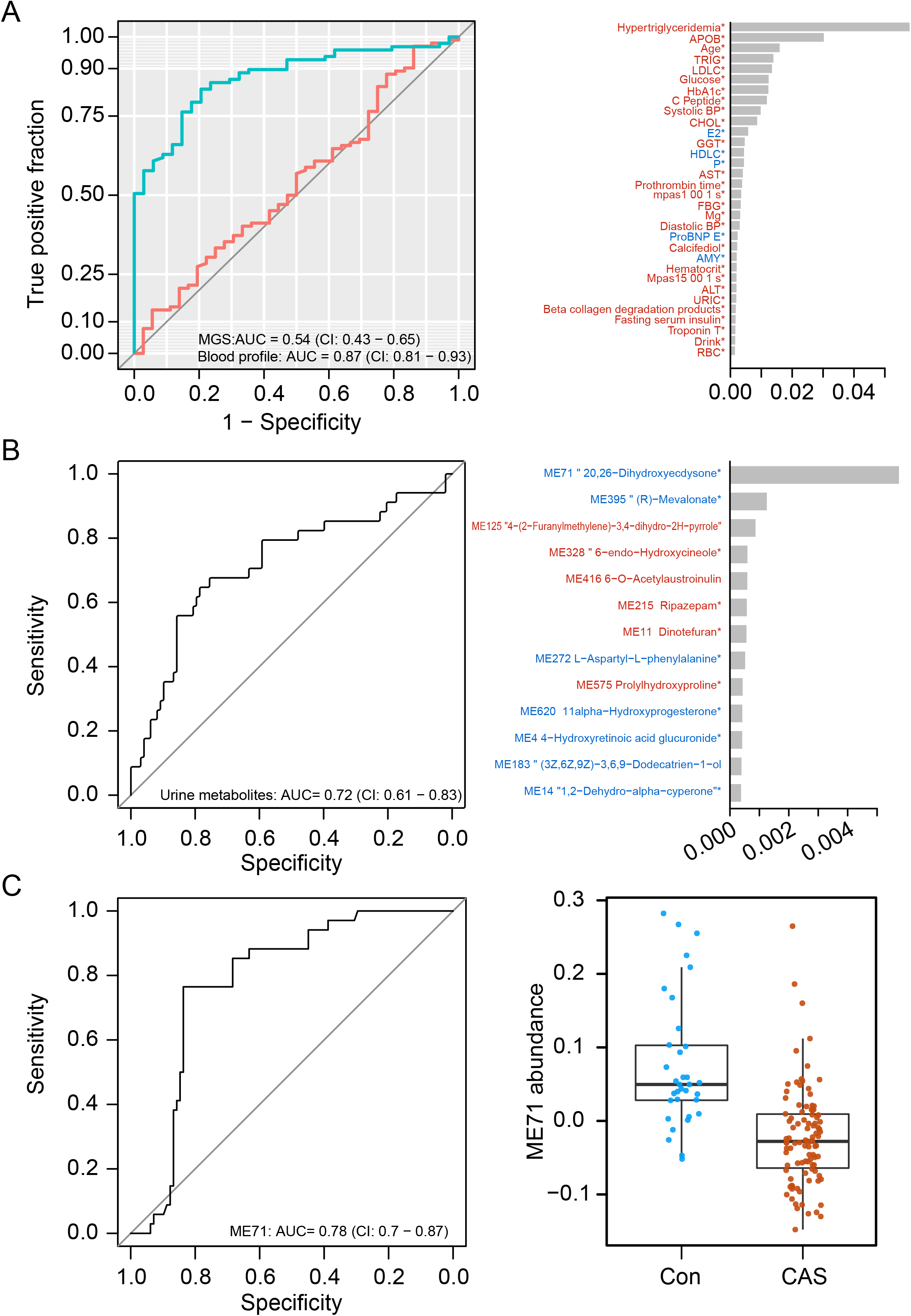
Correlation communities. Community modeling based on connectivity revealed two distinct communities **(a)**, including a large community composed of dense microbe-metabolite correlations **(b)**, and a microbe-metabolite-phenotype triangular correlated community **(c)**. Edge denotes correlation at q<0.1 and coefficient>0.3. The largest community **(b)** was formed by 1049 edges of which 99.1% were between MGS and metabolites (MGS and metabolites are coded, Additional file 15: Table S10). The second largest community **(c)** was formed by 270 edges where the majority of the three domains were connected. Grey lines, positive correlations; red lines, negative correlations.

### Towards identification of carotid atherosclerosis biomarkers

Carotid atherosclerosis is usually asymptomatic with high health risks, and an easily accessible biomarker for this disease is lacking [12]. In our cohort, the carotid atherosclerosis cases were identified based on ultrasound examination of the carotid arteries. We first compared the clinical parameters in groups with or without carotid atherosclerosis. A number of variables differed significantly between individuals with and without carotid atherosclerosis (Additional file 16: Fig S6; and Additional file 17: Table S11). PCoA of the gut microbiota composition failed to distinguish between the microbiota of individuals with and without carotid atherosclerosis (Additional file 18: Fig S7), and the same was seen for microbial gene counts and α-diversity (Additional file 19A: Fig S8A). Mann–Whitney U test identified a few MGS including *Prevotella copri* and *Prevotella bivia* that differed in abundance between individuals with and without carotid atherosclerosis (p<0.05), but none of them passed the false discovery rate of q<0.1 (Additional file 19B: Fig S8B). After gene mapping to both KEGG modules and GMM [13], we found several functions altered in the relative gene abundance in the microbiota of carotid atherosclerosis individuals (Additional file 20: Fig S9). These included the reduction in the abundance of genes involved in methanogenesis and the enrichment of genes involved in the metabolism of low molecular mass carbohydrates in carotid atherosclerosis (Additional file 20: Fig S9). We also attempted to pinpoint the carotid atherosclerosis-related microbial features using a classification model. However, the 5-repeat 10-fold cross-validation random forest model performed better on blood biomarkers (AUC=0.87) than on MGS composition (AUC=0.54, Fig. 6A), indicating a more marked deviation associated with carotid atherosclerosis in the blood profile than in the microbiota. However, since the blood-based discriminators (such as hypertriglyceridemia, APOB, TRIG, LDLC, and HbA1c) are common denominators for metabolic diseases, these are poor biomarkers for carotid atherosclerosis on its own. These results suggested that the gut microbiota in carotid atherosclerosis subjects exhibited only minor alterations, and is not useful on its own as a disease biomarker.

**Fig. 6.**
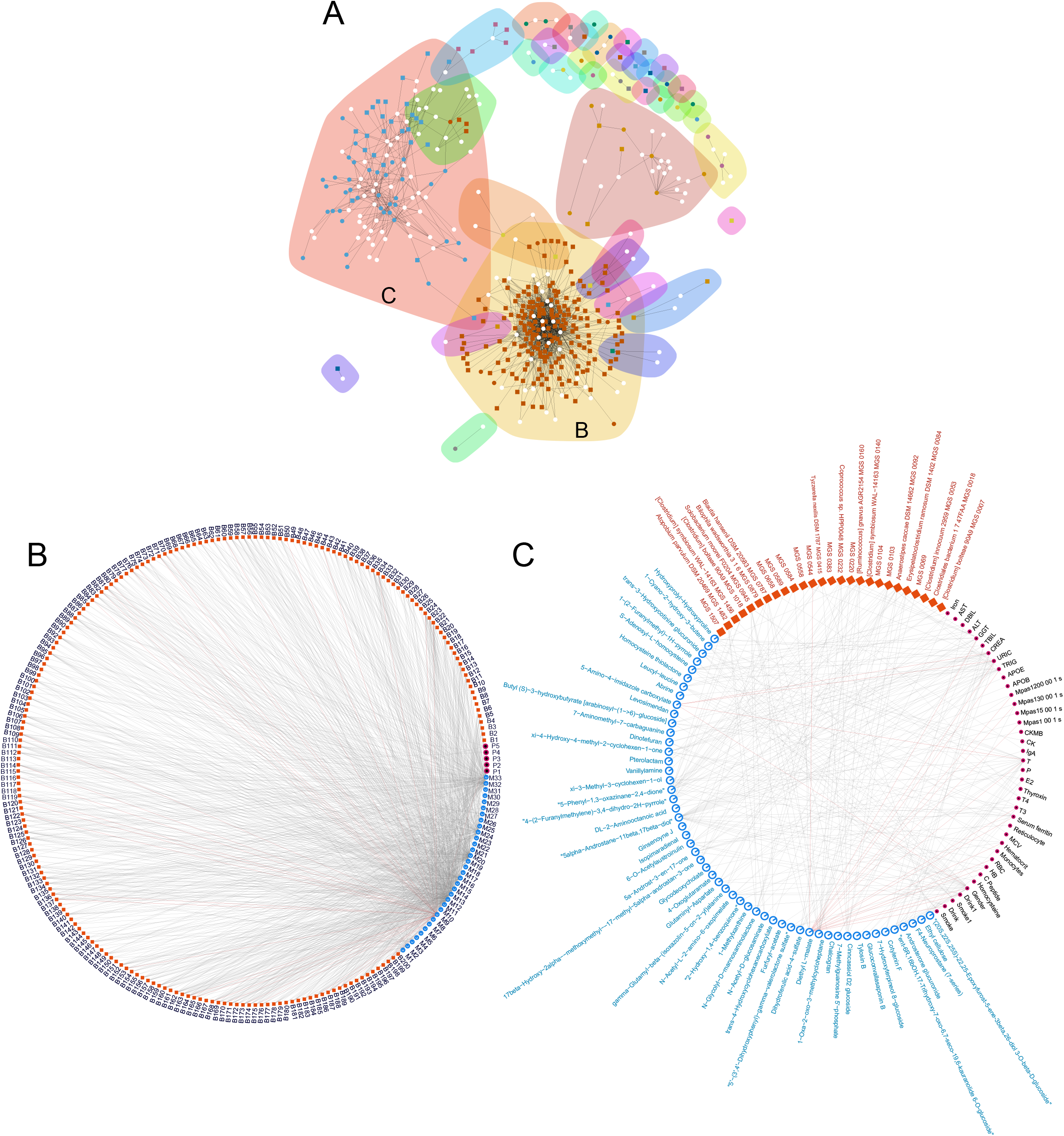
Classification of carotid atherosclerosis based on the gut microbiome or on the urine metabolome. **(a)** ROC curves of carotid atherosclerosis classification based on MGS relative abundances (red) and on the blood profile (green, the left panel), and the top important clinical parameters contributory to the prediction based on blood profile (the right panel). **(b)** The ROC curve of carotid atherosclerosis classification based on the urine metabolome (left panel), and the top important metabolite groups in the classification model (right panel). **(c)** The ROC curve of carotid atherosclerosis classification based on one urine metabolite ME71 (the left panel), and abundances of ME71 in the non-carotid atherosclerosis and carotid atherosclerosis groups (the right panel).

Urine has also previously been used an easily accessible biomarker for several diseases [14]. In search of potential urine biomarkers for carotid atherosclerosis, we likewise conducted feature selection using the 5-repeat 10-fold cross-validation random forest. The model classified individuals with and without carotid atherosclerosis with an AUC of 0.72 based on selected urine biomarkers (Fig. 6B). Notably, the mere inclusion of the top important metabolite group ME71 (structurally similar to 20, 26-dihydroxyecdysone) in the classification model reached an AUC of 0.78 (Fig. 6C, the left panel). This group of metabolites was significantly lower in individuals with vs without carotid atherosclerosis (Fig. 6C, the right panel). However, the variation of ME71 within each group suggests the need for further validation of its use as a biomarker for carotid atherosclerosis.

## Discussion

In the context of the diverse clinical phenotypes in our cohort, we integrated multi-omics data by characterizing inter-domain correlation relationships and further modeled the correlations into communities. The results showed close interactions between the blood profile, the urine metabolome and the gut microbiota, especially that the urine metabolome and the gut microbiota appeared to covary.

One prerequisite of an integrative trans-omics study is that there must be variations in the data points. The metabolic variations between subjects in this study ensured the variation of clinical variables and thereby the related omics, hence providing the necessary preconditions for an integrative study. In addition, the treatment-naïve nature of this cohort eliminated confounding by medication.

Our study, like many previous studies, indicates that changes in the abundance of specific bacteria correlate with fluctuations in certain clinical parameters. These bacteria may serve as putative disease biomarkers and provide potential therapeutic targets. Consistent with findings in our study, the genus *Eubacterium* has been found to be enriched in healthy controls as compared to patients with liver cirrhosis [15], and the abundance of *Ruminococcus* has been associated with liver fibrosis [16]. In mouse models, the administration of *Faecalibacterium prausnitzii* has been shown to improve liver health and reduce hepatic fat accumulation [17]. In comparison to the gut microbiota, our results suggest that alterations of the urine metabolome may greatly co-vary with clinical parameters, and certain groups of urine metabolites may be generally linked to health or disease. These co-varying urine metabolites may be produced or regulated by the same or related metabolic pathways or by specific bacteria.

By conduction a trans-omic network analysis of all three data sets, we provide a view into the pattern of connections between the blood parameters, the gut microbiome, and the urine metabolome. The statistical correlation between the clinical parameters, the gut microbiome, and the urine metabolome does not permit distinction between direct biological crosstalk or co-occurrence. Nonetheless, association relationships may allow for generation of hypotheses to be tested in further studies focusing on mechanisms or etiology. For instance, the metabolic characteristics of certain bacteria (for example, the health-related bacterium *Eubacterium eligens* and the disease-related *Ruminococcus gnavus* are both degraders of complex carbohydrates [18, 19]) is one point worthy of further exploration for the potential mechanisms, and they may leave metabolite fingerprints in the blood and also in the urine that have functional impacts on the host (for example, the well-documented metabolic effects of short-chain fatty acids [20]). The two large communities uncovered by network modeling demonstrate that trans-system relationships are quite clustered and packed, suggesting that entities in the three systems interact frequently. The largest community highlighted the close association between the gut microbiome and the urine metabolome, and therefore the urine metabolomic profile at least in part may reflect the gut microbiota composition.

In a previous study, potential microbial biomarkers for atherosclerotic cardiovascular disease were identified based on 218 cases and 187 controls [10]. However, our study indicates that carotid atherosclerosis is only associated with minor alterations of the gut microbiota. Further studies are needed to identify possible associations between the gut microbiota and carotid atherosclerosis, with adjustment of confounding factors and a larger sample size. Notably, our findings suggest that the urine metabolome may contain promising biomarkers for carotid atherosclerosis, and the potential of the metabolite (ME71) as a carotid atherosclerosis marker warrants further validation.

## Conclusions

Our results provide evidence that changes in clinical features related to disease can be manifested as quantitative changes in multiple body systems including the blood profile, the gut microbiome, and the urine metabolome. These multiple systems are intertwined into closely linked networks. The precise definition and accurate classification of health and disease entail a data-driven, multi-omics and integrative approach for ultimate identification of the most consistent and reliable biomarkers for a given disease.

## Methods

### Study cohort and sample collection

138 Chinese individuals (aged 32-76, including 76 males and 57 females) living in the north of China were recruited during physical examination in a carotid atherosclerosis screening program at the Chinese PLA General Hospital. None of them had been diagnosed with diseases prior to the inclusion, and subjects using antibiotics or receiving other medication within 1 month were excluded. Ultrasound examination of the carotid arteries was used to diagnose carotid arteriosclerosis. Blood tests on 130 clinical parameters were performed. Fecal and urine samples were collected on the day of physical examination, immediately frozen and stored at −80°C until being further processed.

The study was approved by the Medical Ethical Review Committee of the Chinese PLA General Hospital and the Institutional Review Board at BGI-Shenzhen. Informed written consent was obtained from all participants.

### Metagenomic sequencing, binning and annotation

DNA was extracted from fecal samples using Qiagen QIAamp DNA Stool Mini Kit (Qiagen, Germany) according to the manufacturer’s instructions. Construction of a paired-end library with insert size of 350bp was performed according to the manufacturer’s instruction, and the DNA library was sequenced with PE reads of 2×100bp on the Illumina Hiseq2000 platform (Illumina, US). After quality control and host gene removal, clean reads were mapped to the 9,879,896 genes in the integrated gene catalogue [21] with a threshold of more than 90% identity over 95% of the length. Gene abundances were determined as previously described [21], and subsequently used to calculate the relative abundances of genera and KOs. The genes were clustered into MGS based on co-abundance and gene content (≥700 genes) [9]. Taxonomy assignment was based on >50% of the genes in one MGS[5]. The relative abundance of a MGS was calculated from the mean relative abundance of its constituent genes [5]. MGS present in >10% samples were subjected to further analysis.

### Correlation analysis between the gut microbiome, the urine metabolome and blood biomarkers

Spearman correlation was determined between the microbiome (relative abundance of MGS or functions) and blood biomarkers, between the urine metabolome (metabolite groups) and blood biomarkers, and between the urine metabolome and the gut microbiome using the R package. MGS/functions, metabolite groups, and biomarkers that reached statistical significance at three rounds of correlations were retained (p>0.05), and correlations with q<0.1 are indicated in the heatmap.

### Community analysis

Pair-wise Spearman correlation was calculated between MGSs and urine metabolites, between urine metabolites and blood biomarkers, and between MGS and blood parameters. Multiple hypothesis testing was adjusted using the method of Benjamini and Hochberg (q<0.1), and correlations were further filtered by a cutoff of |ρ|≥0.3. Only inter-omic correlations were used for community analysis. Non-directional communities composed of nodes from and edges between the three sets of omics data were modeled using the algorithm proposed by Girvan and Newman [11]. This method involved iterative calculation of edge betweenness centrality on a network, which is the number of weighted shortest paths from all vertices to all other vertices that pass over a given edge. After each iteration, the edges with the highest betweenness centrality were removed, and the process was repeated until only individual nodes remained. Community structure was visualized in R.

### Prediction and classification by random forest

Prediction of clinical parameters was performed by a 5-repeated 10-fold down-sampling cross-validation random forest model (R 3.3.0, randomForest 4.6-12 package) using the relative abundance of MGS as input. Prediction accuracy was evaluated by Spearman correlation between the predicted value and the measured value.

Non-carotid atherosclerosis versus carotid atherosclerosis classification was performed by a 5-repeated 10-fold down-sampling cross-validation random forest model (R 3.3.0, randomForest 4.6-12 package) using blood biomarkers or MGSs as input. Five-repeated 10-fold cross-validation is a robust model matric estimation and down-sampling to deal with the unbalanced sample size between groups. ROC curves were plotted using this classifier with R 3.3.0, pROC package.

### Functional analysis of microbiota

Reporter score (Z-score) [22] was calculated for KO modules or the gut metabolic modules (GMMs) [13] to identify the differentially enriched functions comparing non-carotid atherosclerosis and carotid atherosclerosis groups. Modules with | Z-score | ≥1.96 were considered significant.

### Urine metabolomics

Metabolomic profiling of urine samples was performed on a 2777C UPLC system (Waters, UK) coupled to a Xevo G2-XS QTOF mass spectrometer (Waters, UK). Peak picking and compound identification were performed with Progenesis QI (ver2.2), aligned against the HMDB and KEGG databases. The untargeted metabolite data set was subjected to dimension reduction by the cluster of co-abundant metabolites into groups following the methods provided by a previous study using the R package WGCNA [6]. The positively charged and negatively charged metabolites were analyzed together. Signed weighted co-abundance correlations (biweight midcorrelations after log2 transformation) were calculated across all individuals. A scale-free topology criterion was used to choose the soft threshold of β = 13. Clusters were identified with the dynamic hybrid tree-cutting algorithm, using deepSplit of 4 and a minimum cluster size of 3. The profile of each metabolite cluster was summarized by the cluster eigenvector.

## List of abbreviations

MGS: Metagenomic species
PERMANOVA: Permutational multivariate analysis of variance

Refer to Additional file 1 (Table S1) for abbreviations used for phenotypes.

## Declarations

### Consent for publication

Not applicable.

### Availability of data and material

The data generated and/or analyzed during the current study are available in the CNGB Nucleotide

Sequence Archive (CNSA: https://db.cngb.org/cnsa; accession number CNP0000048)

### Authors’ contributions

All authors read and approved the final manuscript. R.L., Z.J., Q.F., J.W., H.Y., X.X., Y.H., H.J., and K.H. conceived and designed the project. R.L., J.X., C.L. and Z.X. recruited subjects, collected samples, and provided metadata. B.T. coordinated the sample processing at BGI. H.J., Q.F. and Z.J. performed laboratory analysis. Z.J., R.F., Y.G., F.L., and H.X. performed the bioinformatic analyses, and prepared figures and tables. H.Z., L.M. S.B. K.K., X.Y., and H.J. interpreted the data and provided comments. X.Y. wrote the manuscript. K.K., S.B., and L.M. discussed data and revised the manuscript. H.J. and K.H. supervised the project.

## Acknowledgements

We gratefully acknowledge colleagues at BGI-Shenzhen for DNA extraction, library construction, sequencing, and discussions.

## Additional file

Additional file 1: **Table S1**. Abbreviations used for phenotypes.

Additional file 2: **Table S2**. Cohort metadata.

Additional file 3: **Table S3**. Distribution of clinical phenotypes.

Additional file 4: **Table S4**. The metagenomic data for each sample.

Additional file 7: **Table S5**. Correlations between MGS and phenotypes.

Additional file 9: **Table S6**. Correlations between modules and phenotypes.

Additional file 11: **Table S7**. Correlations between urine metabolites and phenotypes.

Additional file 12: **Table S8**. Correlations between MGS and phenotypes and between MGS and urine metabolites that fulfil the triangular correlation.

Additional file 14: **Table S9**. Correlations between modules and phenotypes and between modules and urine metabolites that fulfil the triangular correlation.

Additional file 15: **Table S10**. The code table of MGS, metabolites and phenotypes in Fig 5B.

Additional file 17: **Table S11**. Differential phenotypes in the non-carotid atherosclerosis versus carotid atherosclerosis groups.

**Additional file 5: Fig S1.**
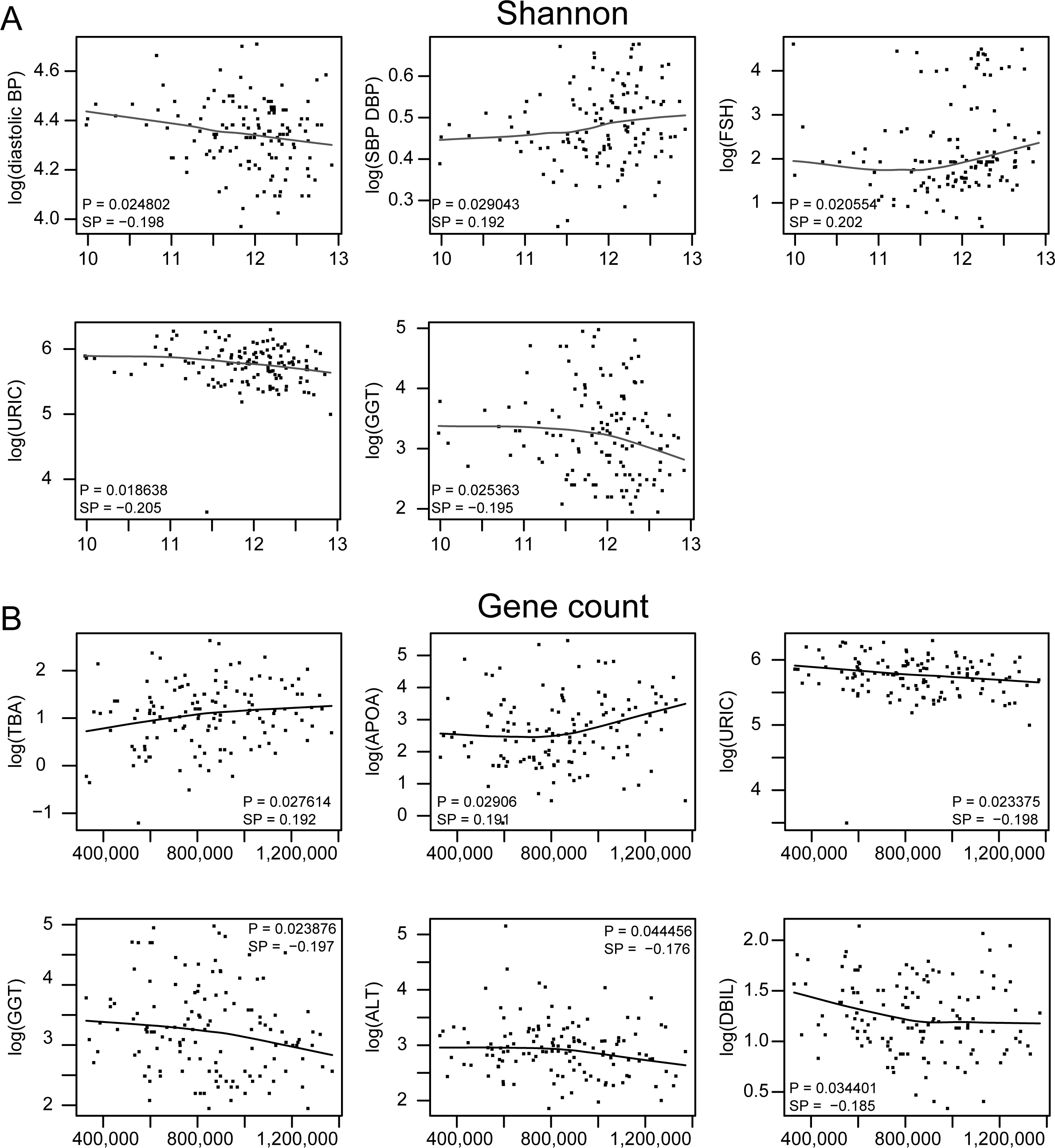
Correlations between Shannon diversity (A) or gene counts (B) with clinical parameters.

**Additional file 6: Fig S2.**
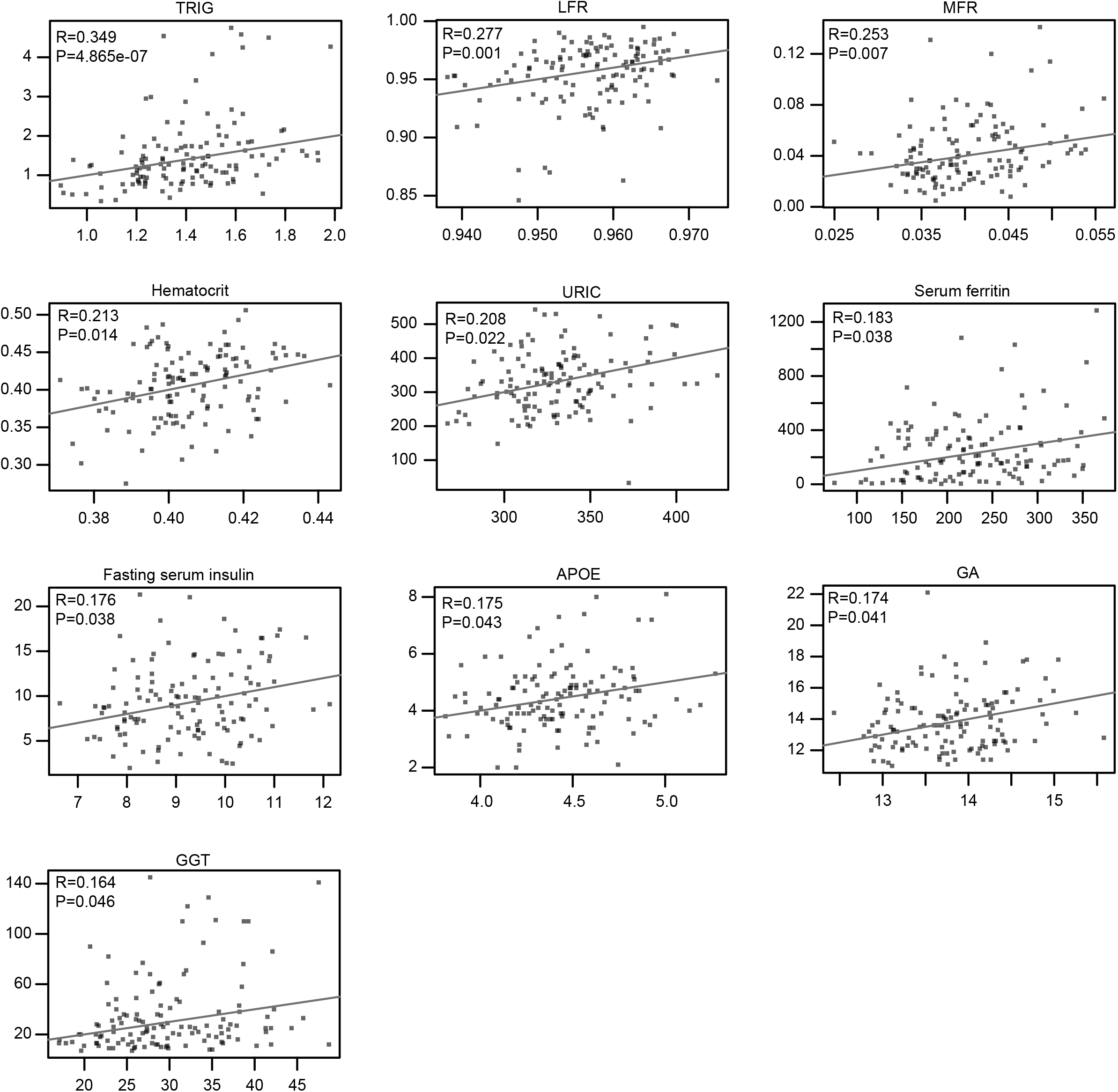
Prediction of clinical parameters by MGS relative abundances with random forest models. Prediction accuracy was evaluated by the correlation between the predicted value and the measured value.

**Additional file 8: Fig S3.**
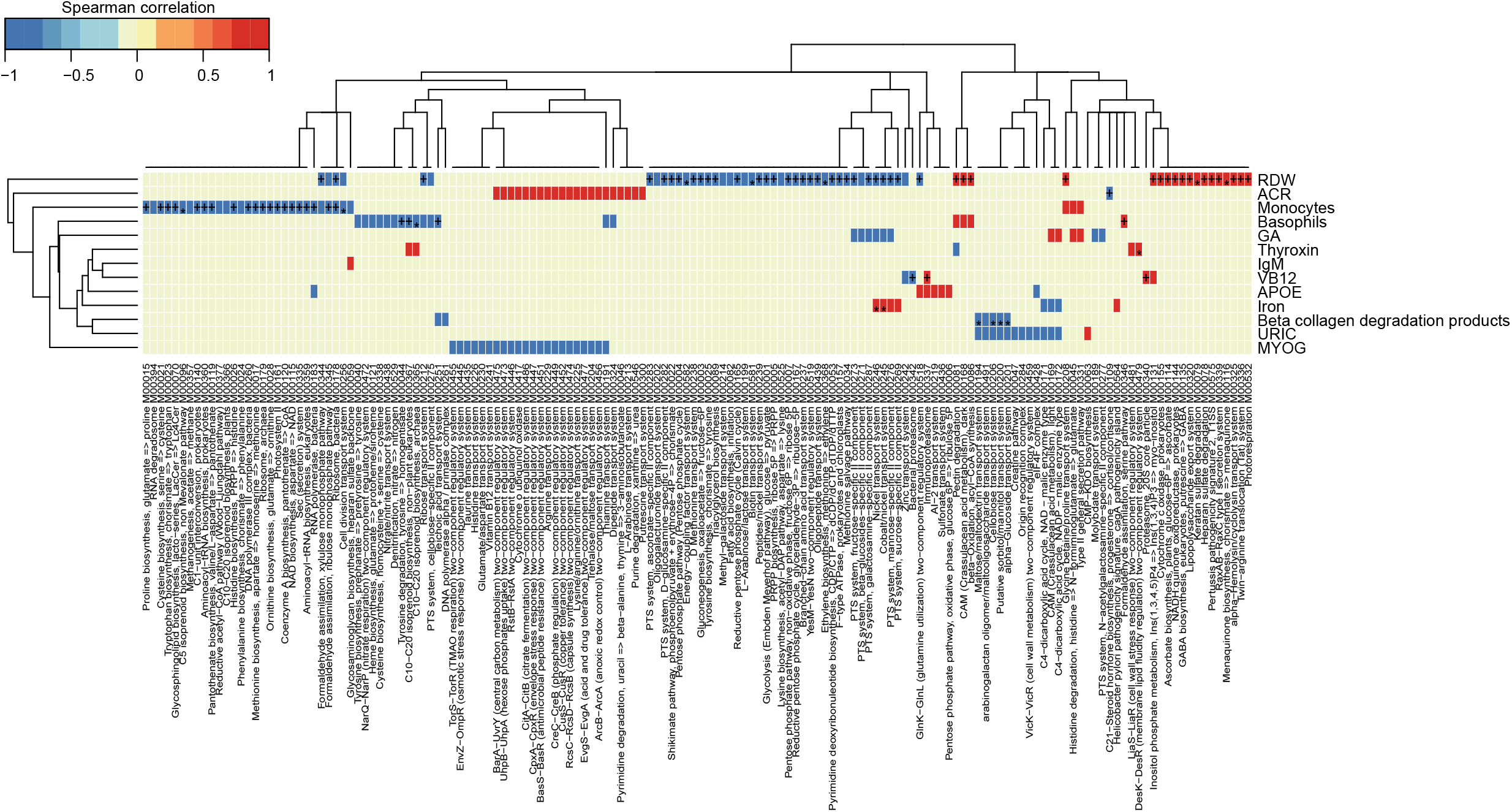
Correlations between microbial functional modules and clinical parameters. Heatmap showing correlations between functional modules and clinical parameters (spearman correlation, p<0.05). +, q<0.1; *, q<0.01. Color scale indicates the value of correlation coefficient.

**Additional file 10: Fig S4.**
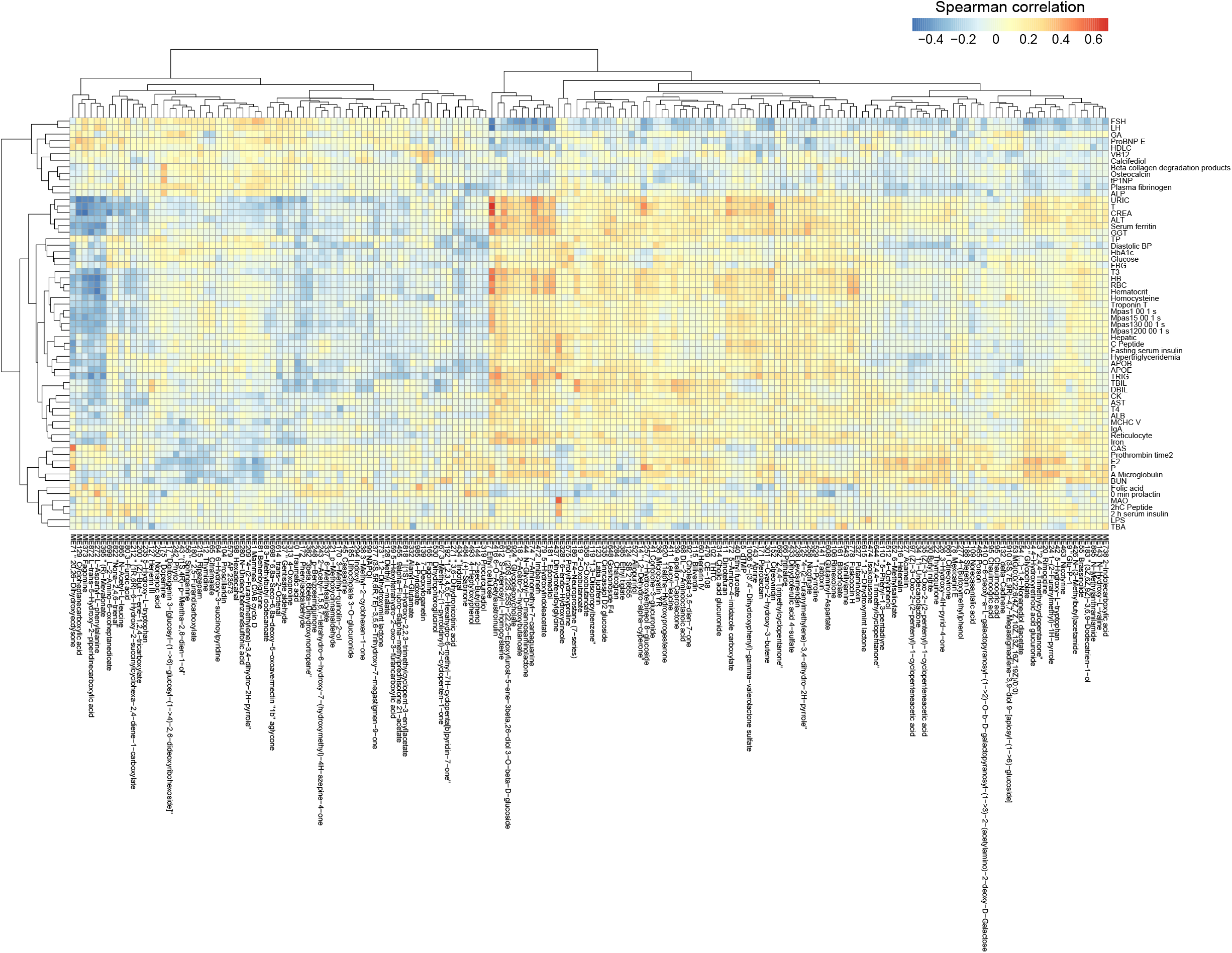
Correlations between urine metabolites and clinical parameters. Heatmap showing all correlations between urine metabolomic groups and clinical parameters (Spearman correlation, p<0.05, q<0.05). Color scale indicates the value of correlation coefficient.

**Additional file 13: Fig S5.**
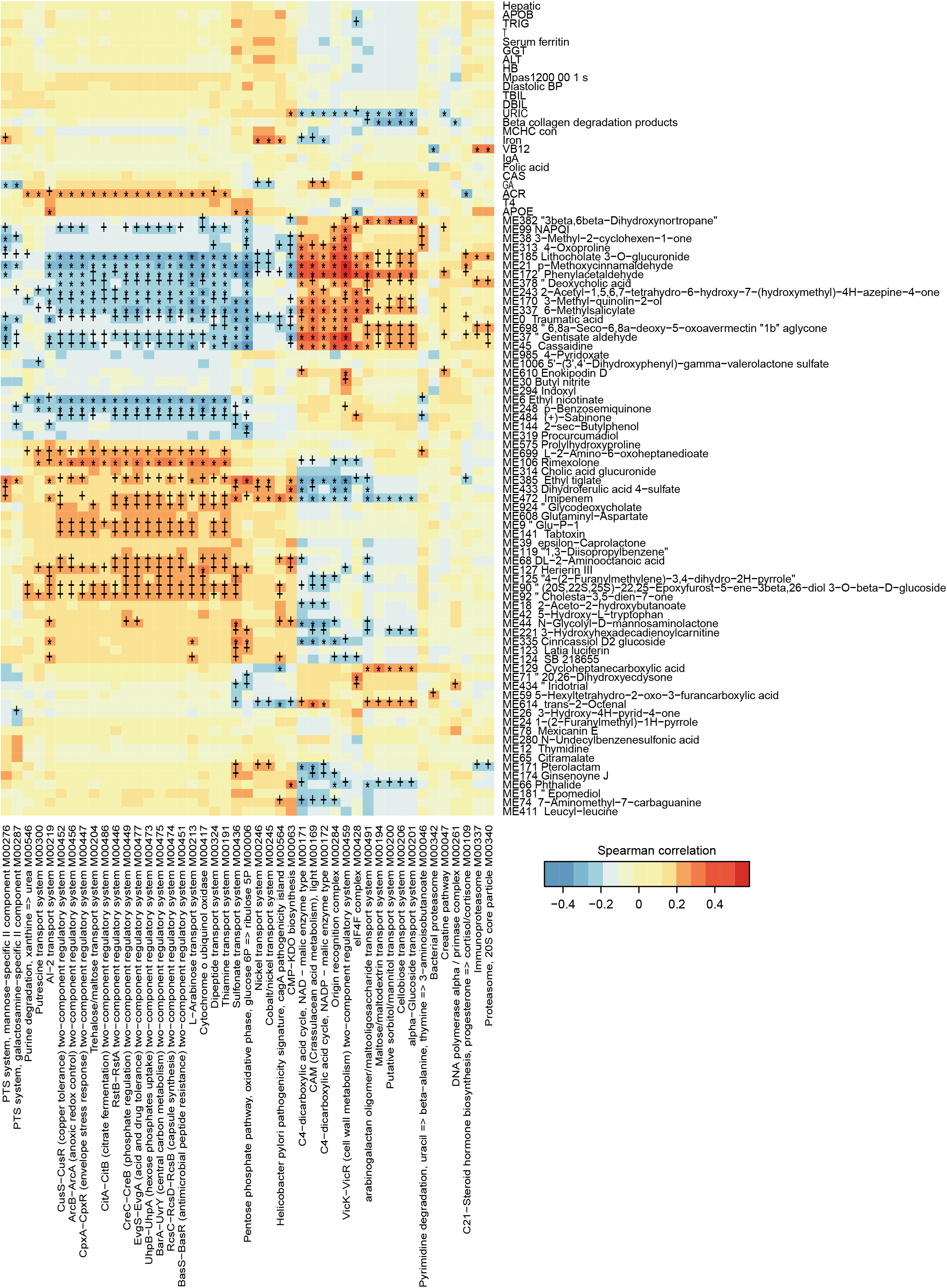
Trans-omics correlations between microbial functional modules, urine metabolites and clinical parameters. Correlations between microbial functions and clinical parameters, between urine metabolites and clinical parameters, and between microbial functions and urine metabolites are shown in the heatmap (spearman correlation, p<0.05). +, q<0.1; *, q<0.01. Color scale indicates the value of correlation coefficient.

**Additional file 16: Fig S6.**
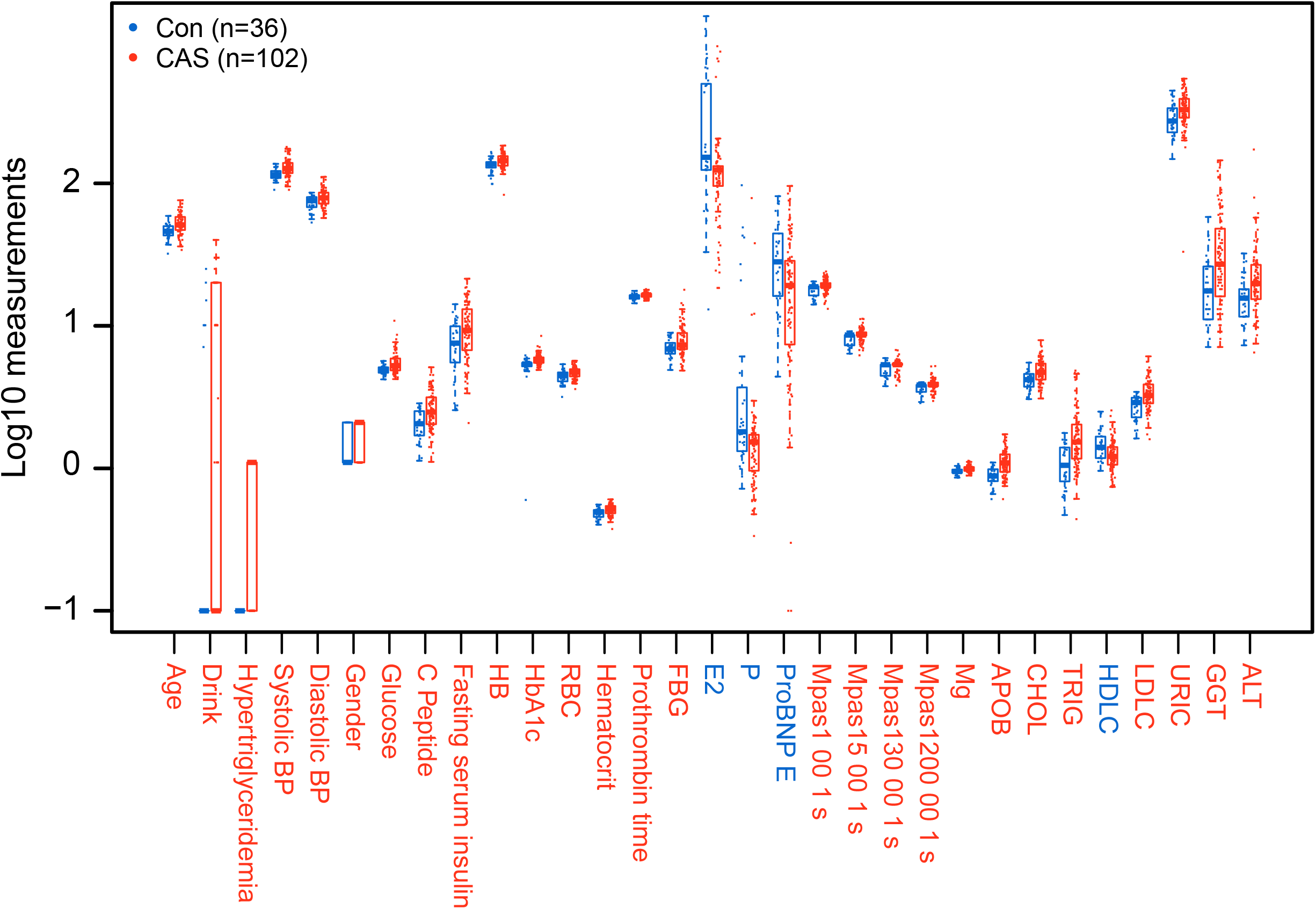
Clinical phenotypes that differed between the non-carotid atherosclerosis and carotid atherosclerosis groups. Wilcoxon rank-sum test was performed between the non-carotid atherosclerosis and carotid atherosclerosis groups.

**Additional file 18: Fig S7.**
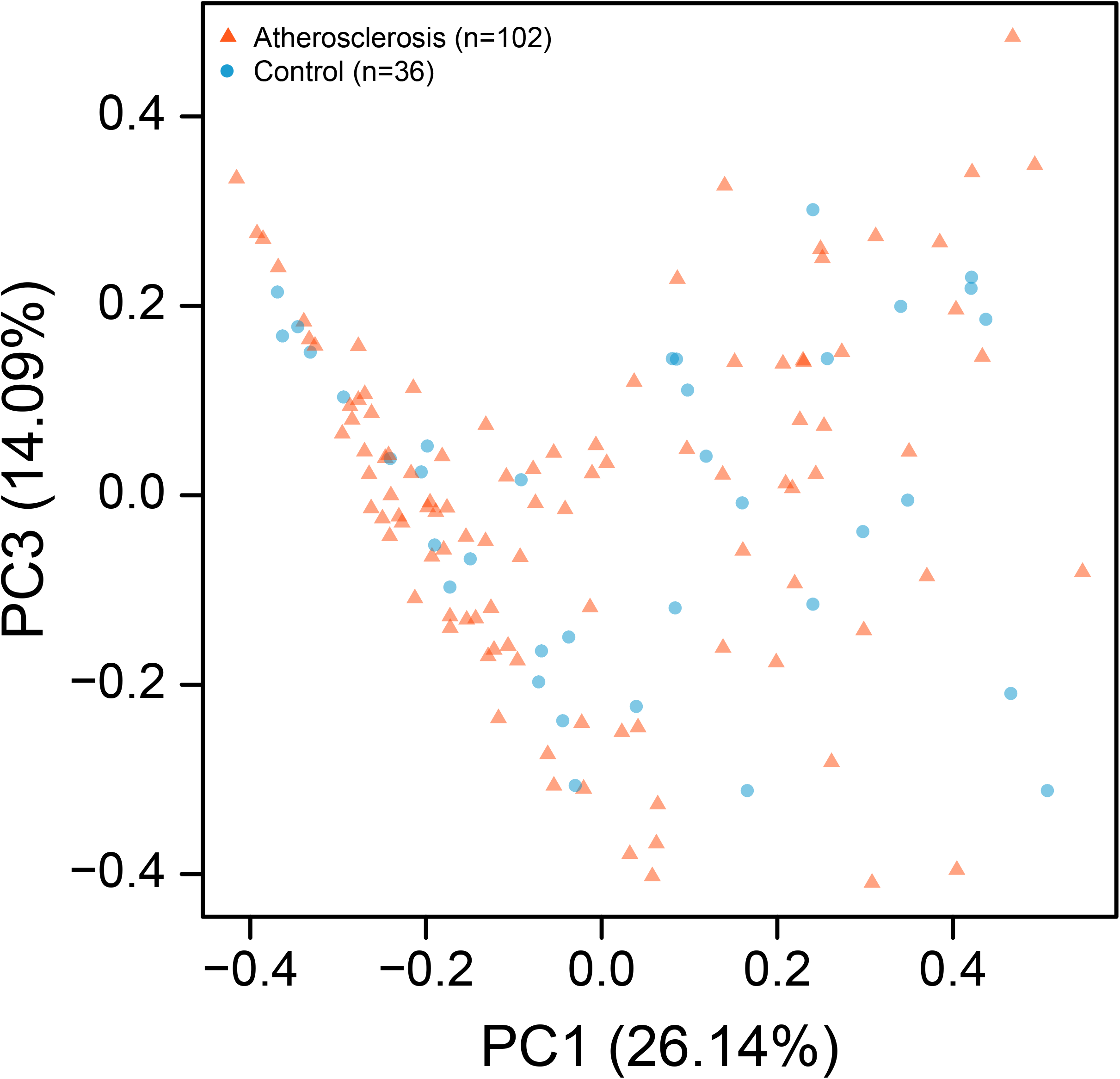
PCoA of genus compositions in the gut microbiota. non-carotid atherosclerosis, blue circles; carotid atherosclerosis, red triangles.

**Additional file 19: Fig S8.**
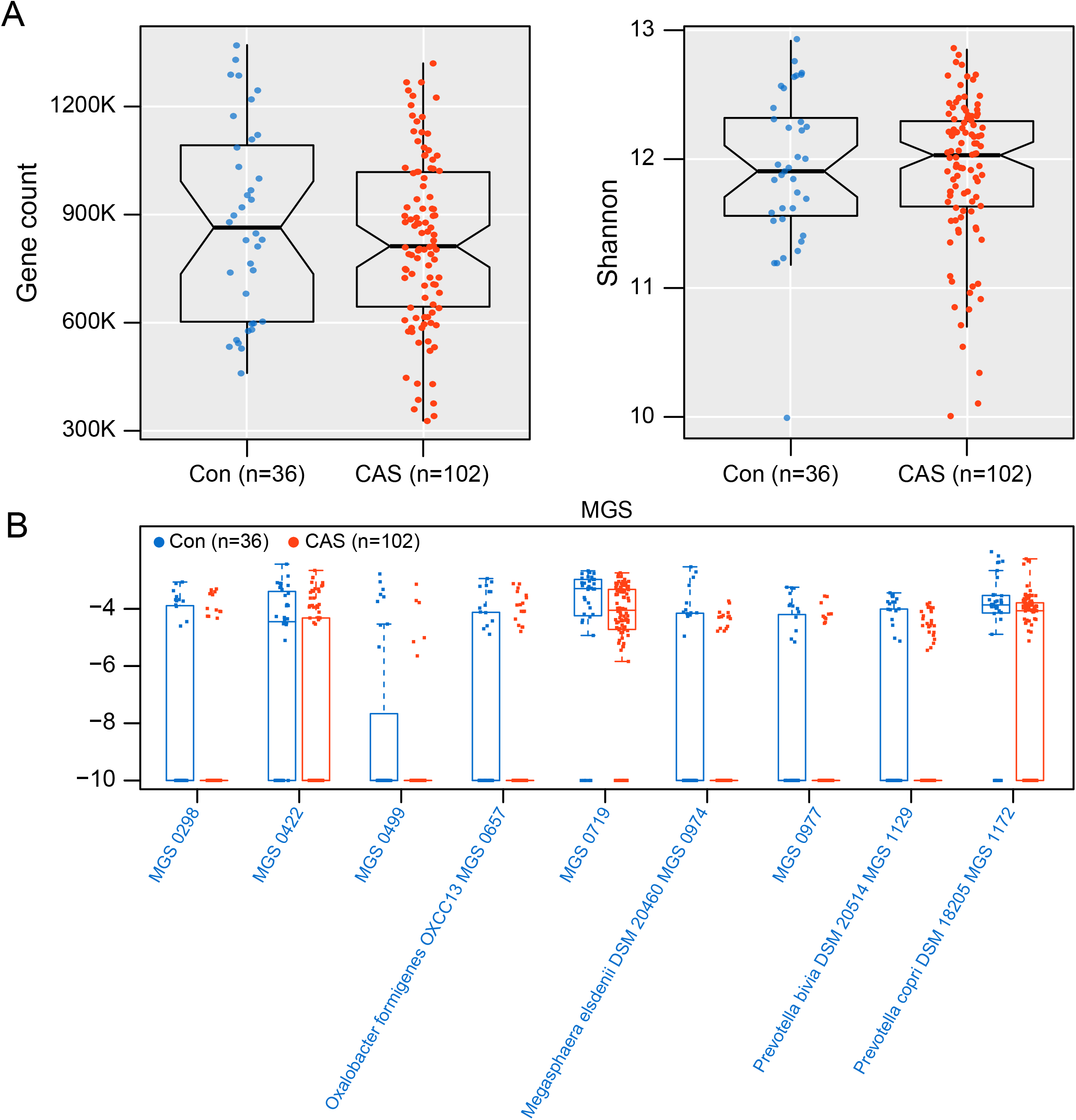
Alterations of the gut microbiota composition in carotid atherosclerosis. (a) Comparison of gene counts (the left panel) and α-diversity (the right panel) of the gut microbiota in non-carotid atherosclerosis (Con) and carotid atherosclerosis individuals (CAS) by Wilcoxon rank-sum test. (b) MGS differing in abundance between non-carotid atherosclerosis and carotid atherosclerosis at p<0.05 but q>0.1.

**Additional file 20: Fig S9.**
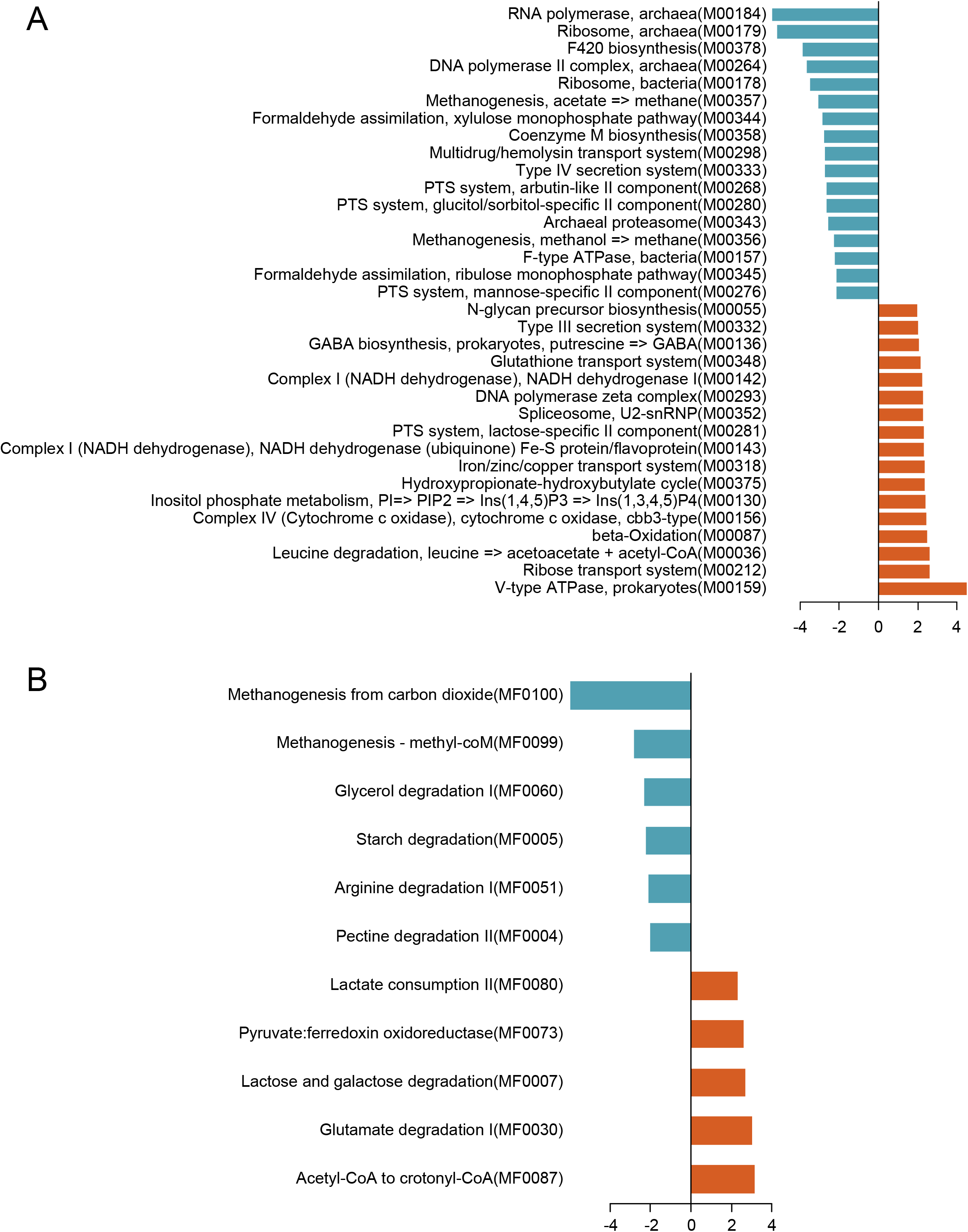
Functional alterations of the gut microbiota in carotid atherosclerosis. Differentially enriched modules analyzed by KEGG (a) and by GMM (b) are shown. Green, enriched in the non-carotid atherosclerosis group; red, enriched in the carotid atherosclerosis group. Bar length indicates the value of reporter score.

